# MAPPING OF LONG-RANGE CHROMATIN INTERACTIONS BY PROXIMITY LIGATION ASSISTED CHIP-SEQ

**DOI:** 10.1101/074294

**Authors:** Rongxin Fang, Miao Yu, Guoqiang Li, Sora Chee, Tristin Liu, Bing Ren

## Abstract

We report a highly sensitive and cost-effective method for genome-wide identification of chromatin interactions in eukaryotic cells. Combining proximity ligation with chromatin immunoprecipitation and sequencing, the method outperforms the state of art approach in sensitivity, accuracy and ease of operation. Application of the method to mouse embryonic stem cells improves mapping of enhancer-promoter interactions.

## Main text

Formation of long-range chromatin interactions is a crucial step in transcriptional activation of target genes by distal enhancers. Mapping of such structural features can help to define target genes for cis regulatory elements and annotate the function of non-coding sequence variants linked to human diseases^1–4^. Study of long-range chromatin interactions and their role in gene regulation has been facilitated by the development of chromatin conformation capture (3C)-based technologies^5,6^. Among the commonly used high-throughput 3C approaches are Hi-C and ChIA-PET^7,8^. Global analysis of long-range chromatin interactions using Hi-C has been achieved at kilobase resolution, but requires billions of sequencing reads^9^. High-resolution analysis of long-range chromatin interactions at selected genomic regions can be attained cost-effectively through either chromatin analysis by paired-end tag sequencing (ChIA-PET), or targeted capture and sequencing of Hi-C libraries^8,10,11^. Specifically, ChIA-PET has been successfully used to study long-range interactions associated with proteins of interest at high-resolution in many cell types and species^12^. However, the requirement for tens to hundreds of million cells as starting materials has limited its application. To reduce the amount of input material without compromising the robustness of long-range chromatin interaction mapping, we developed Proximity Ligation Assisted ChIP-seq (PLAC-seq), which combines formaldehyde crosslinking and *in situ* proximity ligation with chromatin immunoprecipitation and sequencing (**Fig. 1a** and see **Methods**). As detailed below, PLAC-seq can detect long-range chromatin interactions in a more comprehensive and accurate manner while using as few as 100,000 cells, or three orders of magnitude less than published ChIA-PET protocols^8,11^ (**Supplementary Fig. 1a**).

**Figure 1.**
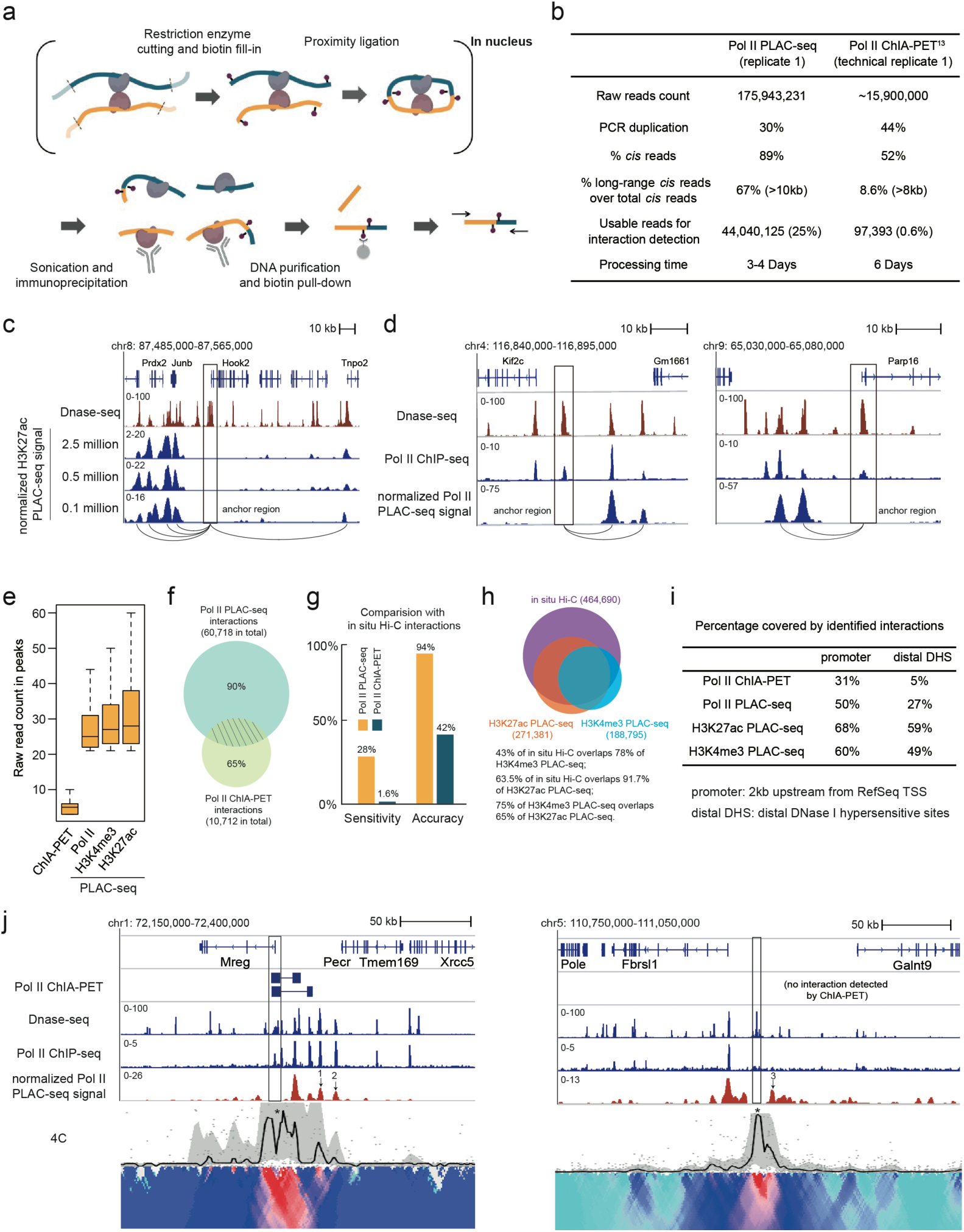
PLAC-seq reveals chromatin interactions in mammalian cells with high sensitivity and accuracy. (a) Overview of PLAC-seq workflow. Formaldehyde-fixed cells are permeabilized and digested with 4-bp cutter MboI, followed by biotin fill-in and *in situ* proximity ligation. Nuclei are then lysed and chromatins, sheared by sonication. The soluble chromatin fraction is then subjected to immunoprecipitation with specific antibodies against either a DNA bound protein or histone modification. Finally, reverse-crosslinking is performed and biotin-labeled ligation junctions are enriched before paired-end sequencing. (**b**) Comparison of sequencing outputs from the Pol II PLAC-seq and ChIA-PET experiments. (**c-d**) Browser plots show examples of high-resolution long-range interactions revealed by H3K27Ac and Pol II PLAC-seq. **c**, promoter-promoter interactions; **d**, left panel, enhancer-enhancer interactions; **d**, right panel, promoter-enhancer interactions. (**e**) Box plots of raw reads count for ChIA-PET and PLAC-seq interactions. (**f**) Overlap between Pol II PLAC-seq and Pol II ChIA-PET interactions. (**g**) Sensitivity and accuracy of PLAC-seq and ChIA-PET interactions compared to *in situ* Hi-C identified interactions. (**h**) Overlap of interactions identified by H3K27ac, H3K4me3 PLAC-seq and *in situ* Hi-C. (**i**) Comparison of coverage of promoters and distal DHSs between PLAC-seq and ChIA-PET. (**j**) Comparison of 4C-seq, PLAC-seq, ChIA-PET anchored at *Mreg* promoter and a putative enhancer (1,2,3 highlight interactions not detected by ChIA-PET; 4C anchor points are marked by asterisk while PLAC-seq and ChIA-PET anchor regions are marked by black rectangle).

We performed PLAC-seq with mouse ES cells and using antibodies against RNA Polymerase II (Pol II), H3K4me3 and H3K37ac to determine long-range chromatin interactions at genomic locations associated with the transcription factor or chromatin marks (**Table 1**). The complexity of the sequencing library generated from PLAC-seq is much higher than ChIA-PET when comparing the Pol II PLAC-seq and ChIA-PET experiments. As a result, we were about to obtain 10x more sequence reads and collecting 440 times more monoclonal *cis* long-range (>10kb) read pairs from a Pol II PLAC-seq experiment than a previously published Pol II ChIA-PET experiment ^13^ (**Fig. 1b**). In addition, PLAC-seq library has substantially fewer inter-chromosomal pairs (11% vs. 48%), but much more long-range intra-chromosomal pairs (67% vs. 9%) and significantly more usable reads for interaction detection (25% vs. 0.6%). Therefore, PLAC-seq is much more cost-effective than ChIA-PET (**Fig. 1b**).

**Table 1.**
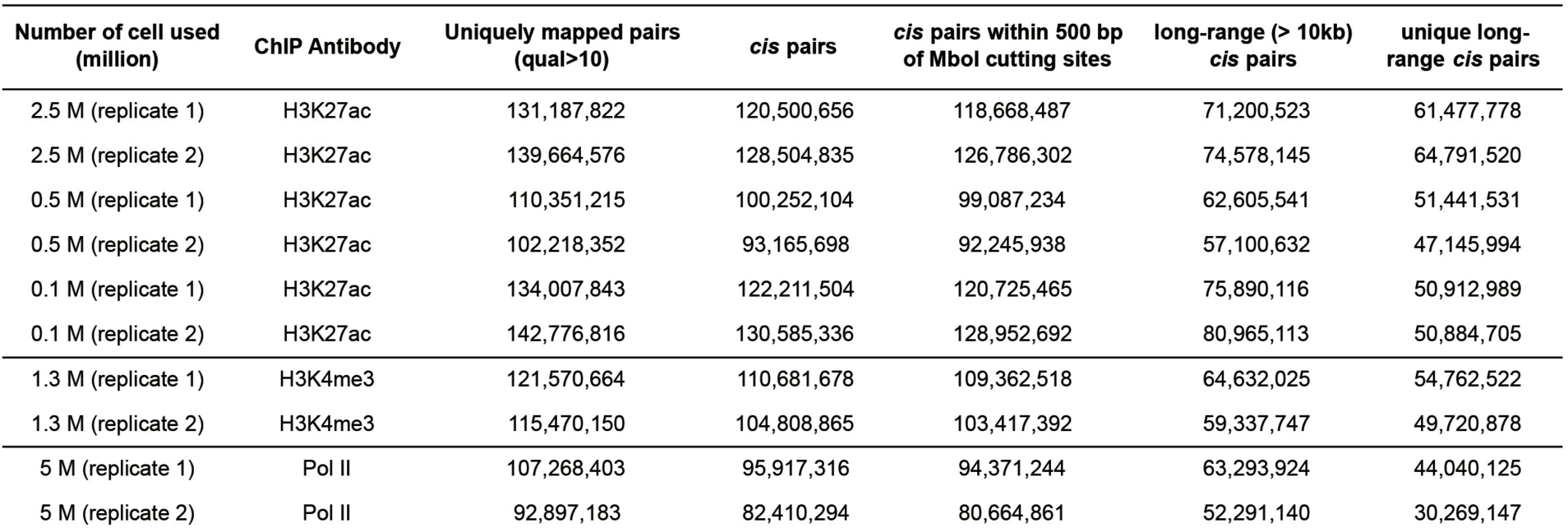
Summary of PLAC-seq libraries

| Number of cell used (million) | ChIP Antibody | Uniquely mapped pairs (qual>10) | *cis* pairs | *cis* pairs within 500 bp of Mbol cutting sites | long-range (> 10kb) *cis* pairs | unique long-range *cis* pairs |
| --- | --- | --- | --- | --- | --- | --- |
| 2.5 M (replicate 1) | H3K27ac | 131,187,822 | 120,500,656 | 118,668,487 | 71,200,523 | 61,477,778 |
| 2.5 M (replicate 2) | H3K27ac | 139,664,576 | 128,504,835 | 126,786,302 | 74,578,145 | 64,791,520 |
| 0.5 M (replicate 1) | H3K27ac | 110,351,215 | 100,252,104 | 99,087,234 | 62,605,541 | 51,441,531 |
| 0.5 M (replicate 2) | H3K27ac | 102,218,352 | 93,165,698 | 92,245,938 | 57,100,632 | 47,145,994 |
| 0.1 M (replicate 1) | H3K27ac | 134,007,843 | 122,211,504 | 120,725,465 | 75,890,116 | 50,912,989 |
| 0.1 M (replicate 2) | H3K27ac | 142,776,816 | 130,585,336 | 128,952,692 | 80,965,113 | 50,884,705 |
| 1.3 M (replicate 1) | H3K4me3 | 121,570,664 | 110,681,678 | 109,362,518 | 64,632,025 | 54,762,522 |
| 1.3 M (replicate 2) | H3K4me3 | 115,470,150 | 104,808,865 | 103,417,392 | 59,337,747 | 49,720,878 |
| 5 M (replicate 1) | Pol II | 107,268,403 | 95,917,316 | 94,371,244 | 63,293,924 | 44,040,125 |
| 5 M (replicate 2) | Pol II | 92,897,183 | 82,410,294 | 80,664,861 | 52,291,140 | 30,269,147 |

To evaluate the quality of PLAC-seq data, we first compared it with the corresponding ChIP-seq data previously collected for mouse ES cells (ENCODE)^14^ and found that PLAC-seq reads were significantly enriched in factor binding sites (*P* < 2.2e-16) and are highly reproducible between biological replicates (Pearson correlation > 0.90) (**Supplementary Fig. 1b-g, Supplementary Fig. 2**). Therefore, we combined the data from two biological replicates for subsequent analysis. We used a published algorithm ‘GOTHiC’^15^ to identify long-range chromatin interactions in each dataset (see **Methods**). We observed highly reproducible interactions identified by H3K27ac PLAC-seq using 2.5, 0.5 and 0.1 million of cells (**Supplementary Fig. 3a**). Furthermore, PLAC-seq signals normalized by *in situ* Hi-C data (see **Methods**) revealed interactions at sub-kilobasepair resolution even with 100,000 cells (**Fig. 1c-d**). We identified a total of 60,718, 271,381, and 188,795 significant long-range interactions from Pol II, H3K27ac or H3K4me3 PLAC-seq experiment, respectively. Previously, ChIA-PET was performed for Pol II in mouse ES cells, providing us a reference dataset for comparison^13^. After examining the raw read counts from the PLAC-seq interacting regions, we found that each chromatin contact was typically supported by 20 to 60 unique reads. By contrast, chromatin interactions identified in ChIA-PET analysis were generally supported by fewer than 10 unique pairs^13^ (**Fig. 1e**). Next, we found that Pol II PLAC-seq analysis identified a lot more interactions than Pol II ChIA-PET (~60,000 vs. ~10,000), with 10% PLAC-seq overlapping with 35% of ChIA-PET intra-chromosomal interactions (FDR < 0.05 and PET count >= 3) (**Fig. 1f**). To further investigate the sensitivity and accuracy of each method, we performed *in situ* Hi-C on the same cell line and collected 300 million unique long-range (> 10kb) *cis* pairs from ~1.2 billion paired-end sequencing reads. Using ‘GOTHiC’ (see **Methods**), 464,690 long-range chromatin interactions were identified. We found that 94% of the chromatin interactions found in Pol II PLAC-seq overlapped with 28% of *in situ* Hi-C interactions (see **Methods**), while 44% of contacts detected by ChIA-PET matched less than 2% of that of in siteu Hi-C contacts (**Fig. 1g**). We also examined H3K27ac and H3K4me3 PLAC-seq interactions and found that the interactions identified by these two marks together recovered 68% of the *in situ* Hi-C interactions (**Fig. 1h**). In addition, we observed that PLAC-seq interactions in general have a higher coverage on regulatory elements such as promoters and distal DNase I hypersensitive sites (DHSs) compared to ChIA-PET (**Fig. 1i**). Taken together, the results above support the superior sensitivity and specificity of PLAC-seq over ChIA-PET. To further validate the reliability of PLAC-seq, we performed 4C-seq analysis at four selected regions (**Supplementary Table 1**). Although most interactions were independently detected by both ChIA-PET and PLAC-seq methods (**Fig. 1j**, left panel, and **Supplementary Fig. 3b**), we found three strong interactions (marked 1,2,3 in **Fig. 1j**) determined by 4C-seq that were detected by PLAC-seq, but not ChIA-PET. Conversely we also identified a case of chromatin interaction uniquely detected by ChIA-PET but not observed from 4C-seq (highlighted by red rectangle in **Supplementary Fig. 3b**), once again supporting the superior performance of PLAC-seq over ChIA-PET.

Next, we focused on H3K4me3 and H3K27ac PLAC-seq datasets to study promoter and active enhancer interactions in the mouse ES cells. As expected, we found that PLAC-seq interactions are highly enriched with the corresponding ChIP-seq peaks compared to *in situ* Hi-C interactions (**Fig. 2a**). The enrichment allowed us to further explore interactions specifically enriched in PLAC-seq compared to *in situ* Hi-C due to chromatin immunoprecipitation. Identifying such interactions may help us understand higher-order chromatin structures associated with a specific protein or histone mark. To achieve this, we developed a computational method (https://github.com/r3fang/PLACseq) using Binomial test to detect interactions that are significantly enriched in PLAC-seq relative to *in situ* Hi-C (see **Methods**). We termed this type of interactions as ‘PLACE’ (<u>PLAC</u>-<u>E</u>nriched) interactions. A total of 28,822 and 19,429 significant H3K4me3 or H3K27ac PLACE interactions (*q* < 0.05) (**Supplementary Table 2,3**) in the mouse ES cells were identified, respectively. 26% of H3K27ac PLACE interactions overlapped with 19% of H3K4me3 PLACE interactions, suggesting that they contain different sets of chromatin interactions (**Fig. 2b**). Indeed, we found majority of H3K27ac PLACE interactions are enhancer-associated interactions (74%) while H3K4me3 PLACE interactions are generally associated with promoters (78%) (**Fig. 2c**). The difference between H3K27ac and H3K4me3 PLACE interactions prompted us to further explore these two types of interactions. We examined the expression level of genes associated with H3K27ac and H3K4me3 PLACE interactions and discovered that genes involved in H3K27ac PLACE interactions have a significantly higher expression level than genes associated with H3K4me3 PLACE interactions (P < 2.2e-16, **Fig. 2d**), suggesting that the former assay could be used to discover chromatin interactions at active enhancers.

**Figure 2.**
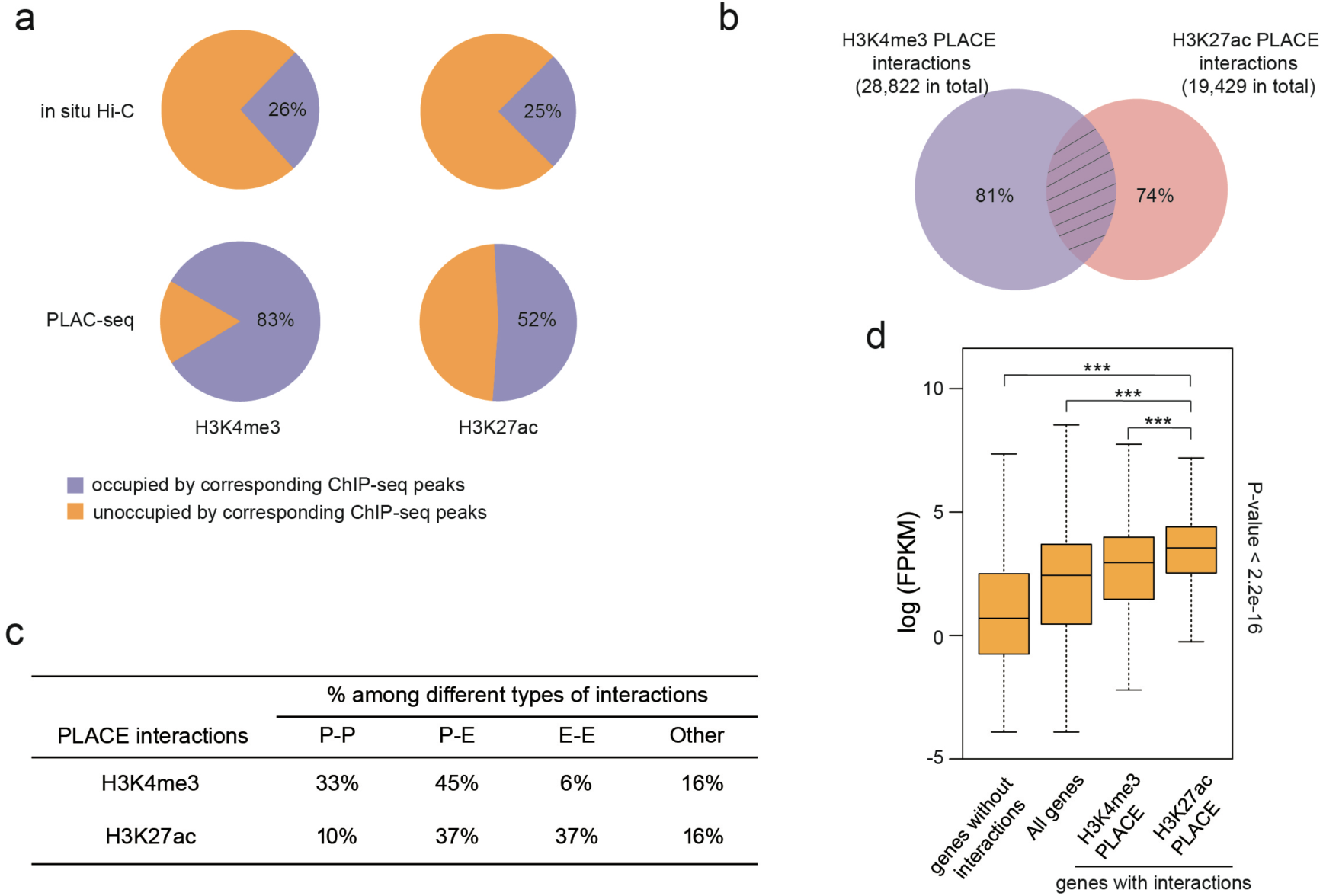
H3K4me3 and H3K27ac PLAC-seq data identify promoter and enhancer interactions in mESC. (**a**) PLAC-seq interactions are enriched at genomic regions associated with the corresponding histone modifications. (**b**) Overlap between H3K27ac and H3K4me3 PLAC-Enriched (PLACE) interactions. (**c**) Distribution of promoter-promoter, promoter-enhancer, enhancer-enhancer and other interactions for H3K27ac and H3K4me3 PLACE interactions. (**d**) Boxplot of expression of different groups of genes. H3K27ac PLACE interactions are associated with genes express significantly higher than other genes (Wilcoxon tests, *P*<2.2e-16).

In summary, we developed a new method to map long-range chromatin interactions in a eukaryotic genome. Our data suggest that PLAC-seq can generate more comprehensive and accurate interaction maps than ChIA-PET. Using PLAC-seq, we obtained an improved map of enhancer and promoter interactions in mouse embryonic stem cells. The ease of experimental procedure, low amount of cells, cost-effectiveness of this method will allow it be broadly adopted, thereby greatly facilitating the mapping of long-range chromatin interactions in a much broader set of species, cell types and experimental settings than previous approaches.

## Acknowledgement

The work is supported by funding from the Ludwig Institute for Cancer Research and NIH (1U54DK107977-01 and U54 HG006997) to B.R.

## Author contribution

MY and RF designed the experiments. MY performed the PLAC-seq experiments. GL and TL carried out the in situ Hi-C and 4C experiments. RF carried out data analysis. RF, MY, SC and BR prepared the manuscript.

## Online Methods

### Cell culture and fixation

The F1 *Mus musculus castaneus* × S129/SvJae mouse ESC line (F123 line) was a gift from the laboratory of Dr. Rudolf Jaenisch and was previously described^16^. F123 cells were cultured as described previously^17^. Cells were passaged once on 0.1% gelatin-coated feeder-free plates before fixation.

To fix the cells, cells were harvested after accutase treatment and suspended in medium without Knockout Serum Replacement at a concentration of 1x10^6^ cells per 1ml. Methanol-free formaldehyde solution was added to the final concentration of 1% (v/v) and rotated at room temperature for 15 min. The reaction was quenched by addition of 2.5 M glycine solution to the final concentration of 0.2 M with rotation at room temperature for 5 min. Cells were pelleted by centrifugation at 3,000 rpm for 5 min at 4 °C and washed with cold PBS once. The washed cells were pelleted again by centrifugation, snap-frozen in liquid nitrogen and stored at -80 °C.

### PLAC-seq protocol

PLAC-seq protocol contains three major parts: *in situ* proximity ligation, chromatin immunoprecipitation or ChIP, biotin pull-down followed by library construction and sequencing. The *in situ* proximity ligation and biotin pull-down procedures were similar to previously published *in situ* Hi-C protocol^9^ with minor modifications as described below:

1. *In situ* proximity ligation. 0.5 to 5 millions of crosslinked F123 cells were thawed on ice, lysed in cold lysis buffer (10 mM Tris, pH 8.0, 10 mM NaCl, 0.2% IGEPAL CA-630 with proteinase inhibitor) for 15 min, followed by a washing step with lysis buffer once. Cells were then resuspended in 50 µl 0.5% of SDS and incubated at 62 °C for 10 min. Permeabilization was quenched by adding 25 µl 10% Triton X-100 and 145 µl water, and incubation at 37 °C for 15 min. After adding NEBuffer 2 to 1x and 100 units of MboI, the digestion was performed for 2 h 37 °C in a thermomixer, shaking at 1,000 rpm. After inactivation of MboI at 62 °C for 20 min, biotin fill-in reaction was performed for 1.5 h 37 °C in a thermomixer after adding 15 nmol of dCTP, dGTP, dTTP, biotin-14-dATP (Thermo Fisher Scientific) each and 40 unit of Klenow. Proximity ligation was performed at room temperature with slow rotation in a total volume of 1.2 ml containing 1xT4 ligase buffer, 0.1 mg/ml BSA, 1% Triton X-100 and 4000 unit of T4 ligase (NEB).
2. ChIP. After proximity ligation, the nuclei were spun down at 2,500 g for 5 min and the supernatant was discarded. The nuclei were then resuspended in 130 µl RIPA buffer (10 mM Tris, pH 8.0, 140 mM NaCl, 1 mM EDTA, 1% Triton X-100, 0.1% SDS, 0.1% sodium deoxycholate) with proteinase inhibitors. The nuclei were lysed on ice for 10 min and then sonicated using Covaris M220 with following setting: power, 75 W; duty factor, 10%; cycle per burst, 200; time, 10 min; temp, 7 °C. After sonication, the samples were cleared by centrifugation at 14,000 rpm for 20 min and supernatant was collected. The clear cell lysate was mixed with Protein G Sepharose beads (GE Healthcare) and then rotated at 4 °C for pre-clearing. After 3h, supernatant was collected and ~5% of lysate was saved as input control. The rest of the lysate was mixed with 2.5 µg of H3K27Ac (ab4729, abcam), H3K4me3 (04-745, millipore) or 5 µg Pol II (ab817, abcam) specific antibody and incubate at 4 °C overnight. On the next day, 0.5% BSA-blocked Protein G Sepharose beads (prepared one day ahead) were added and rotated for another 3 h at 4 °C. The beads were collected by centrifugation at 2,000 rpm for 1 min and then washed with RIPA buffer three times, high-salt RIPA buffer (10 mM Tris, pH 8.0, 300 mM NaCl, 1 mM EDTA, 1% Triton X-100, 0.1% SDS, 0.1% sodium deoxycholate) twice, LiCl buffer (10 mM Tris, pH 8.0, 250 mM LiCl, 1 mM EDTA, 0.5% IGEPAL CA-630, 0.1% sodium deoxycholate) once, TE buffer (10 mM Tris, pH 8.0, 0.1 mM EDTA) twice. Washed beads were first treated with 10 µg Rnase A in extraction buffer (10 mM Tris, pH 8.0, 350 mM NaCl, 0.1 mM EDTA, 1% SDS) for 1 h at 37 °C. Then 20 µg proteinase K was added and reverse crosslinking was performed overnight at 65 ^o^C. The fragmented DNA was purified by Phenol/Chloroform/Isoamyl Alcohol (25:24:1) extraction and ethanol precipitation.
3. Biotin pull-down and library construction. The biotin pull-down was performed according to *in situ* Hi-C protocol with the following modifications: 1) 20 µl of Dynabeads MyOne Streptavidin T1 beads were used per sample instead of 150 µl per sample; 2) To maximize the PLAC-seq library complexity, the minimal number of PCR cycles for library amplification was determined by qPCR.

### PLAC-seq and Hi-C read mapping

We developed a bioinformatics pipeline (https://github.com/r3fang/PLACseq) to map PLAC-seq and in-situ Hi-C data. We first mapped paired-end sequences using BWA-MEM^18^ to the reference genome (mm9) in single-end mode with default setting for each of the two ends separately. Next, we paired up independently mapped ends and only kept pairs if each of both ends were uniquely mapped (MQAL>10). As we focused only on intrachromosomal analysis in this study, interchromosomal pairs were discarded. Next, read pairs were further discarded if either end was mapped more than 500bp apart away from the closest MboI site. Read pairs were next sorted based on genomic coordinates followed by PCR duplicate removal using MarkDuplicates in Picard tools. Finally, the mapped pairs were partitioned into “long-range” and “short-range” if its insert size was greater than the given distance of default threshold 10kb or smaller than 1kb, respectively.

### PLAC-seq visualization

For each given anchor point, we first extracted the interaction read pairs with one end falling in the anchor region, the other flanking outside it. Next, we focused on the 2MB window surrounding the anchor point and split this region into a set of 500bp non-overlapping bins. We extended the flanking read into 2kb, then counted the coverage for each bin from both PLAC-seq and *in situ* Hi-C experiments. The read count was later normalized into RPM (Read Per Million) and the final normalized PLAC-seq signal was the subtraction between treatment and input.

### PLAC-seq and *in situ* Hi-C interaction identification

We used ‘GOTHiC’^15^ to identify long-range chromatin interactions in PLAC-seq and *in situ* Hi-C datasets with 5kb resolution. To identify the most convincing interactions, we consider an interaction significant if its FDR < 1e-20 and read count > 20. In total, we identified 60,718, 271,381, 188,795 significant long-range interactions from Pol II, H3K27ac, H3K4me3 PLAC-seq and 464,690 from *in situ* Hi-C in the mouse ES cells.

### Interaction overlap

We define that two distinct interactions are overlapped if both ends of each interaction intersect by at least one base pair.

### Identification of PLACE interactions

H3K4me3/H3K27ac/Pol2 ChIP-seq peaks in mouse ES cells were downloaded from ENCODE^14^. We expanded each peak to 5kb as an anchor point. PLAC-Enriched (PLACE) interactions can be identified by the exact binomial test using *in situ* Hi-C as an estimation of background interaction frequency. In greater detail, for each anchor region *i*, we first counted the number of read pairs have one end overlap with anchor region *read_total_total*_*i*_ and *read_total_total*_*i*_ for PLAC-seq and *in situ* Hi-C. Next, we focused on a 2MB window flanking the anchor and partitioned this region into a set of overlapping 5kb bins with a step size of 2.5kb. Briefly, the probability that a read pair is the result of a spurious ligation between the anchor region *i* and bin *j* can be estimated as:

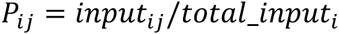

Then, the probability of observing *treat*_*i,j*_ read-pairs in PLAC-seq between *i* and bin *j* can be calculated by the binomial density:

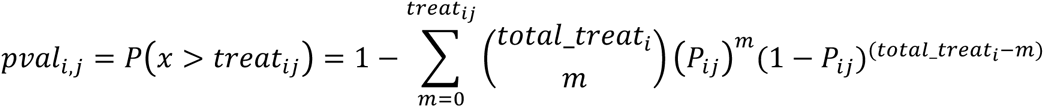

Next, bins that have a binomial *P* value smaller than 1e-5 were identified as candidates. Centering on each candidate, we chose a 1kb, 2kb, 3kb, 4kb window and calculated the fold change respectively, then defined the peak with the largest fold change as an interaction:

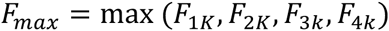

Overlapping interactions were merged as one interaction and binomial *P* was recalculated based on the merged interaction. Next, the resulting *P* values were corrected to *q* value to account for multiple hypothesis testing using Bonferroni correction. Finally, interactions with *q* value smaller than 0.05 are reported as significant interactions.

### Hi-C and PLAC-seq contact maps visualization

*In situ* Hi-C or PLAC-seq contact maps were visualized using Juicebox^19^ after removing all *trans* reads and *cis* reads pairs span less than 10kb.

### 4C validation

4C experiments were performed as previously described^20^. The restriction enzymes used and the primer sequences for PCR amplification are listed in **Supplementary Table 1**. Data analysis was performed using 4Cseqpipe^21^.

### *In situ* Hi-C

F123 *in situ* Hi-C was performed as previously described with 5 million of F123 cells^9^.

